# Sexual selection, environmental robustness and evolutionary demography of maladapted populations: a test using experimental evolution in seed beetles

**DOI:** 10.1101/426056

**Authors:** Ivain Martinossi-Allibert, Emma Thilliez, Göran Arnqvist, David Berger

## Abstract

Whether sexual selection impedes or aids adaptation has become a pressing question in times of rapid environmental change and parallels the debate about how the evolution of individual traits impacts on population dynamics and viability. The net effect of sexual selection on population viability results from a balance between genetic benefits of “good genes” effects and costs of sexual conflict. Depending on how these facets of sexual selection are affected under environmental change, extinction of maladapted populations could either be avoided or accelerated. Here, we evolved seed beetles under three alternative mating regimes (polygamy, monogamy and male-limited selection) to disentangle the contributions of sexual selection, fecundity selection and male-female coevolution to individual reproductive success and population fitness. We compared these contributions between the ancestral environment and two new stressful environments (temperature and host plant shift). Evolution under polygamy resulted in the highest individual reproductive success in competitive context for both sexes. Moreover, females evolving only via sexual selection on their male siblings in the male-limited regime had similar reproductive success and higher fertility than females evolving under monogamy, suggesting that sexual selection on males had positive effects on female fitness components. Interestingly, male-limited sexual selection resulted in males that were robust to stress, compared to males from the two evolution regimes applying fecundity selection. We quantified the population-level consequences of this sex-specific adaptation and found evidence that costs of socio-sexual interactions were higher in male-limited lines compared to polygamous lines, and that this difference was particularly pronounced at elevated temperature to which males from the male-limited regime were more robust compared to their conspecific females. These results illustrate the tension between individual-level adaptation and population-level viability in sexually reproducing species and suggest that sex-specific selection can cause differences in environmental robustness that may impact population demography under environmental change.

## Introduction

Evolutionary rescue critically depends on that genetic responses are rapid enough to allow populations to track changes in their environment while the demographic cost of maladaptation remains small enough to avoid genetic drift and extinction (Bell and Gonzales 2009, Walters et al. 2012, Gonzales et al. 2013, Carlson et al. 2014, Orr and Unckless 2014). Research has highlighted the potential discrepancy between adaptation in individual traits and that of the population as a whole (e.g. Cam et al. 2002, Violle et al. 2007, Bolnick et al. 2011, Rankin et al. 2011). This discrepancy may have fundamental influence on the potential for evolutionary rescue because those strategies that maximize individual fitness may often lead to overexploitation of ecological resources, and therefore do not always translate into maximal population viability (Hardin 1968, Kokko and Brooks 2003, Rankin and Lopez-Sepulcre 2005). In sexually reproducing species, these dynamics can become of particular importance, because while population growth is ultimately limited by female egg production, adaptation in traits increasing male fertilization success may evolve irrespective of their effects on population viability (Clutton-Brock and Parker 1995, Arnqvist and Rowe 2005, Rankin et al. 2011). This realization has sparked considerable debate about whether sexual selection generally will act to increase or decrease extinction risk (e.g. Zahavi 1975; Hamilton and Zuk 1982; Manning 1984; Maynard-Smith 1991; Lorch et al. 2003; Kokko and Brooks 2003; Whitlock and Agrawal 2009; Rankin et al. 2011; Holman and Kokko 2014; Gerber and Kokko 2016; Martínez-Ruiz and Knell 2017; De Lisle et al. 2018; Li and Holman 2018; Martinossi-Allibert et al. 2018a, 2018b).

Sexual selection in polygamous species often acts through pre- and post-copulatory female mate choice or male-male competition based on morphological or behavioral traits such as mating calls, antlers, body size, coloration, or sperm characteristics (Andersson 1994). Hypotheses suggesting that sexual selection should increase population fitness assume that the expression and maintenance of these traits is energetically costly and therefore reflects the bearer’s overall condition and genetic quality (Zahavi 1975; Hamilton and Zuk 1982; Jennions *et al*. 2001). Strong sexual selection could therefore target large parts of the genome and purge deleterious mutations with pleiotropic effects on survival and female fecundity (the genic capture hypothesis, Rowe and Houle 1996; Tomkins *et al*. 2004). Moreover, it would do so while leaving females relatively spared of the cost of adaptation, allowing population fitness and viability to remain largely unaffected in the process (Manning 1984; Agrawal and Whitlock 2009).

However, this idea has been contested by studies on a variety of systems that have revealed misalignment of selection in the sexes (*reviewed in*: Bonduriansky and Chenoweth 2009; Rice and Gavrilets 2014). One fundamental consequence of this is that, because males and females share most of their genome, alleles favored in one sex may be detrimental when expressed in the other, resulting in a genetic conflict known as intralocus sexual conflict (IaSC: Rice and Chippindale 2001). IaSC may thus reduce or even reverse any positive contribution of good genes sexual selection to population viability. Sexual selection can also have direct detrimental effects on the population level. This can happen if the successful male strategy inflicts harm on the female during the mating interaction or mate guarding, reducing her fecundity or longevity. Indeed, such male strategies have been observed in a wide range of animal taxa (Parker 1979; Thornhill and Alcock 1983; Clutton-Brock and Parker 1995; Arnqvist and Rowe 2005; Parker 2006). This form of conflict, referred to as interlocus sexual conflict (IeSC), thus represents a type of “tragedy of the commons”, in which male adaptations can in theory drive a population to extinction by overexploiting the main resource limiting population growth (the female and her egg production) (Kokko and Brooks 2003; Rankin *et al.* 2011).

These costs and benefits of sexual selection may produce variable outcomes at the population level, as observed in experimental systems providing evidence for sexual selection increasing adaptation (e.g. Fricke and Arnqvist 2007; Mallet *et al.* 2011; Mcguigan *et al.* 2011; Plesnar-Bielak *et al.* 2012; Sharp and Agrawal 2013; Grieshop *et al.* 2016) as well as impeding it (e.g. Holland 2002; Rundle *et al.* 2006; Hollis and Houle 2011; Arbuthnott and Rundle 2012; Chenoweth *et al.* 2015, Berger et al. 2016a). The impact of environmental change on these dynamics has started to be explored in recent years (e.g. Plesnar-Bielak *et al.* 2012; Arbuthnott et al. 2014; Berger et al. 2014; Connallon and Clark 2014; Punzalan et al. 2014; Gerber and Kokko 2016; Holman and Jacomb 2017; Gomez-Llano et al. 2018; Martinossi-Allibert et al. 2018a; 2018b; Parrett and Knell 2018; Skwierzynska et al. 2018; Yun et al. 2018), and there are indeed reasons to suspect that the different facets of sexual selection will be sensitive to rapid ecological change. For example, male reproductive success often exhibits genotype-by-environment interactions (GEI:s) (Bussiere *et al.* 2008; Kolluru 2014; Miller and Svensson 2014). Such pronounced changes in genotype-ranking across environments may simply reflect strong sexual selection favoring locally adapted male genotypes (e.g. Martinossi-Allibert et al. 2017), but also bring up the question of whether sexual selection generally favors genotypes of high environmental robustness. If sexually selected traits are costly, as predicted by theory, sexual selection could favor resource allocation decisions that under certain circumstances lead to increased mortality in the population (e.g. Brooks 2000; Hunt *et al.* 2004; Kim and Velando 2016; Zajitschek and Connallon 2017). On the other hand, if secondary sexual traits are honest signals of condition (Zahavi 1975; Hamilton and Zuk 1982; Jennions *et al.* 2001), good genes sexual selection may instead favor high quality genotypes that are resilient to most types of stress. While this question has received attention in studies measuring GEI:s for traits presumably involved in sexual selection (*reviewed in*: Bussiere *et al.* 2008; Kolluru 2014; Miller and Svensson 2014), experiments directly linking sexual selection to the evolution of GEI:s and resilience to rapid environmental change are scarce. Moreover, an underappreciated consequence of sex-specificity in such GEI:s are potential knock-on effects at the population level; because the extent of male-induced harm on females should depend on the frequencies of alternative male mating strategies and the relative condition of the mean male and female phenotype (Clutton-Brock and Parker 1995; Parker 2006; Rankin et al. 2011; Takahashi et al. 2014; Iglesias-Carrasco *et al.* 2018), IeSC may be modulated in accordance with the sensitivity of each sex to the change ecological conditions. However, this last hypothesis remains largely unexplored.

Here we contrasted the contribution of sexual selection to population fitness in well-adapted populations assayed in their ancestral environment, and when reared on a new host plant or at elevated temperature, to which they were maladapted. To do this we first subjected replicate lines of seed beetle to experimental evolution under three alternative mating system regimes: Polygamy (allowing sexual selection, fecundity selection, and male-female coevolution), enforced Monogamy (allowing only fecundity selection and minimizing sexual conflict), and Male-limited selection (allowing only sexual selection on males and prohibiting females to coevolve with males). Following 16-20 generations of experimental evolution, the lines were reared in the ancestral or the novel environments and males and females were assayed for their individual reproductive success in competitive settings. At the ancestral and elevated temperature we also assayed the beetles’ fertility as monogamous pairs and their joint offspring production in larger groups. These assays allowed us to explore how evolution under alternative mating systems and levels of sexual versus fecundity selection affected individual-level and population-level estimates of fitness, as well as the link between them, in ancestral and novel environments. It also allowed us to quantify IeSC in terms of the net cost of mating interactions. Specifically, we could infer the importance of sexual selection for female fitness by comparing offspring production in male-limited and polygamous females evolving with sexual selection, to that in monogamous females evolving without it. We could also assess how the opportunity for male-female coevolution affected the impact of sexual conflict on population viability by comparing offspring production in groups of beetles from the polygamous lines (allowing male-female coevolution) and male-limited lines (where female counter-adaptation was prevented).

Evolution with sexual selection conferred high fertility and individual reproductive success in both sexes across environments, in line with “good genes” effects. However, there was also a substantial cost of mating (IeSC), which counteracted the increase in fertility associated with sexual selection. Moreover, this cost was significantly lower in the Polygamous relative to the Male-limited regime, highlighting the importance of male-female coevolution in mitigating the impact of IeSC. In the Male-limited regime, where only sexual selection had been operating, males showed greater environmental robustness to both host plant and temperature compared to the Polygamy and Monogamy regime, suggesting that the fecundity selection included in the two latter regimes had favored alleles encoding male environmental sensitivity. Strikingly, IeSC seemed to be modulated by this sex-specificity: IeSC tended to be reduced at elevated temperature in the Polygamous and Monogamous evolution regimes, but maintained at high level in the Male-limited evolution regime. Our results thus demonstrate that (i) sexual selection can lead to evolutionary increases in female fitness components across environments, but (ii) can also favor male adaptations that are harmful to females and increase IeSC, and (iii), provide proof-of-concept for the idea that sex-specific selection can generate sex-specificity in environmental robustness which in turn can modulate sexual conflict and demography of maladapted populations.

## Methods

### Study Population

The seed beetle *Callosobruchus maculatus* is a common pest on fabaceaous seeds usually found in tropical and subtropical regions where it commonly infests seed storages. Larvae develop inside the beans of their host and emerge as reproductively mature adult; during adult life, they do not require food or water to complete their life-cycle (Fox *et al.* 2011). One of the natural environments of *C. maculatus*, seed storages in tropical regions, is easy to reproduce in the laboratory, which makes it an ideal model system (Fox *et al.* 2003; Messina and Jones 2009). All the beetle stocks that were used in this study were maintained under controlled temperature (29°C), humidity (50%RH) and light cycle (12L: 12D), and reared on the preferred host plants *Vigna unguiculata* (black-eyed bean). Under these conditions, adult lifespan lays typically between 7 to 10 days. The base population that was used in the present study is a conglomerate of 41 isofemale lines that were isolated from a natural population sampled in Lomé, Togo (06°10#N 01 °13#E) in 2010 (see Berger et al. 2014). Creating isofemale lines from the original population allowed to maintain under laboratory conditions a snapshot of the genetic variation present in that population at the time of sampling (Hoffmann and Parsons 1988), which was then pooled together at the start of the experiment. Prior to the start of the experiments and during the experimental evolution protocol, isofemale lines were maintained under controlled temperature (29°C), humidity (50%RH) and light cycle (12L: 12D), and reared on its preferred host plant *Vigna unguiculata* (black-eyed bean). The base populations was created by mixing 30 randomly selected individuals from each isofemale line. After two generations the large base population (N > 3000) was split into three replicate populations. Each replicate was then split among three evolution regimes that were maintained for 16 generations prior to the first experiment (see below).

### Evolution regimes

To decouple the effects of fecundity selection, sexual selection, and male-female coevolution on individual reproductive success and population fitness, we allowed beetles to evolve under three alternative mating regimes: Polygamy (allowing both fecundity and sexual selection, and male-female coevolution), Monogamy (allowing only fecundity selection on male-female mating pairs), and Male-limited selection (allowing only sexual selection on males and preventing male-female coevolution). Effective population size in each regime was kept approximately equal (Ne = 150) by first estimating the variance in lifetime reproductive success expected for each sex, based on previously published data on this population (e.g. Berger et al. 2014, 2016b; Martinossi-Allibert et al. 2017), and then using it to estimate the population size necessary to obtain a Ne of 150 following the equation provided in Falconer and MacKay (1996):

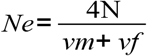

where *vm* and *vf* are represent respectively the variance in reproductive success of males and females.

The number of beans provided as egg laying substrate in each regime was standardized to give the same, relatively low juvenile density (2-4 eggs/bean), to minimize (and equalize) larval competition.

#### Polygamy

Under the Polygamy regime, both males and females experienced selection and had opportunities to mate multiply; this evolution regime was close to natural conditions or regular laboratory maintenance conditions. Each generation, 300 adult seed beetles were transferred to a glass jar containing approximately 4800 black-eyed beans. During 48 hours, individuals were free to interact, copulate and lay eggs on the available beans. After 48 hours, adults were removed from the jar and the beans were saved until emergence of the next generation at which point 300 individuals were randomly collected to seed the new generation.

#### Monogamy

Under the Monogamy regime, virgin individuals were paired at random in monogamous couples in order to remove sexual selection. 0-72h post adult emergence, 123 couples were created and each couple was left to interact during 5 hours in a 6cm petri dish, to allow for multiple matings and male-female interactions. After that, all females were gathered in a glass jar containing approximately 4800 black-eyed beans and left 48 hours to lay eggs. After 48 hours, females were removed and the beans were saved until the emergence of the next generation. In this regime, selection should have been acting on female fecundity and ability to oviposit on high quality substrate (by selecting larger beans or beans presenting low competition, free of eggs laid by other females), as well as on males with positive effects on the fecundity of their female via ejaculate composition and mating behavior.

#### Male-limited selection

Under the Male-limited regime, selection on females was removed while sexual selection was allowed to act in males. 100 virgin individuals of each sex were placed in a 1L glass jar containing a cardboard structure but no beans. This provided a more complex environment than a simple empty jar and allowed individuals to find hiding places (which they normally find among the beans) without having to provide beans on which females would have laid eggs. After 48 hours, during which individuals interacted and copulated at will, females were removed from the jar and placed in individual 6cm petri dishes containing ca. 30 black-eyed beans, where they were left for 48 hours to lay eggs. The next generation was formed by collecting one random offspring of each sex from each dish. This insured that very weak selection was acting in females because all had the same genetic contribution to the next generation, except for the few females (a maximum of three in any generation) that died before egg laying, whereas sexual selection will have favored the males fertilizing the highest fraction of female eggs.

One of the replicates of the Male-limited evolution regime was lost due to mishandling during the experimental evolution protocol, which brought the number of replicates used in the present study to three for the Polygamy and Monogamy regimes and two for the Male evolution regime.

### Experimental design

Competitive lifetime reproductive success (LRS) of individual males and females was measured after 16 generations of experimental evolution, followed by one generation of relaxed selection. Fertility following a single monogamous mating, “population fitness” of mixed-sex groups, and traits putatively related to reproductive success (body weight, ejaculate weight, locomotor activity) were measured after 20 generations of experimental evolution, and one subsequent generation of relaxed selection. The environmental sensitivity of competitive LRS of the lines was assayed in individuals raised as larvae in one of three environments; the ancestral condition (29°C with black-eyed beans *Vigna ungulata* as a larval host), and two stressful conditions; elevated temperature (36°C, black-eyed beans) and host plant shift (adzuki beans *Vigna angularis* at 29°C). Fertility and population fitness were assayed only at the ancestral and elevated temperature conditions due to logistic limitations. The environmental sensitivity of LRS could be estimated independently in males and females by competing them against a reference stock population raised at the ancestral conditions. Fertility and population fitness assays, on the other hand, estimated offspring production for each line as a whole. Moreover, from these latter two estimates, we could quantify the cost of mating interactions (i.e. level of IeSC), by comparing the per female offspring production in the two assays.

#### Competitive LRS

Thirty mating pairs were created from each replicate line of the evolution regimes and split equally among the three larval environments, resulting in 10 pairs per treatment. These pairs were then allowed to mate and produce offspring. Between 5 and 10 offspring per sex and mating pair were scored for competitive LRS, for a total of 3672 assays evenly distributed across evolution regimes, sexes and environments.

Competitive LRS was measured by competing focal males and females from the evolution regimes to a reference population formed some 60 generations previously from the same genetic stock, kept at the same abiotic lab conditions as the evolution regimes under the natural polygamous mating regime. A virgin focal individual (raised in one of the three larval environments) was placed in a 9cm petri dish containing a non-limiting amount of black-eyed beans, together with two virgin individuals of the opposite sex from the reference population, as well as one sterilized competitor of the same sex from the reference population. Importantly, all reference individuals were raised in the ancestral environment (29°C on black-eyed beans), so that putative developmental sensitivity to the novel environments could be attributed solely to the sex and evolution regime of the focal individual. Hence, all assays of competitive LRS had to be performed in the ancestral environment in the adult stage. The presence of a sterilized competitor ensured that focal individuals competed for mating opportunities, as well as post-mating fertilization success in the case of males, and egg laying sites in the case of females, while all emerging offspring in an assay could be attributed to the focal individual (Eady 1991; Maklakov and Arnqvist 2009; Grieshop *et al.* 2016; Martinossi-Allibert *et al.* 2017). Sterilization was achieved by exposing reference individuals to gamma radiation at a dose of 100Gy, which leaves them sterile for their entire lifetime, while leaving no noticeable effects on competitiveness (e.g. Grieshop et al. 2016). Individuals were left to interact during their entire lifetime. After emergence of all offspring of the next generation (following 35 days since the start of the assay), petri dishes were frozen at -20°C for at least two days before the offspring were counted.

#### Fertility, phenotypes and population fitness

These assays were performed on two replicate lines from each evolution regime, maintained at the two temperature conditions. Duplicates were made of each of the 6 lines and split among the two temperatures. Virgin individuals were collected from each line and used in fertility assays and population fitness assays (20 replicates per assay type, line and temperature). Here, as we were aiming to score the temperature sensitivity of each evolution regime as a whole (a combined estimate for conspecific males and females), in monogamous single pair settings (fertility) and multiple mating population settings, both assay types could be carried out at respective temperature and did not have to be limited to juvenile development. Body weight, male ejaculate weight and locomotor activity were measured for the individuals that were used in monogamous assays in order to estimate covariation between fertility and phenotypes. We note that female body weight and male locomotor activity have previously been shown to predict fecundity and the level of sexual conflict in the stock population (Berger et al. 2016a, 2016b)

Fertility was measured by counting the lifetime adult offspring production of a female after a single monogamous mating with a randomly assigned male from her own population. After the mating event, the male and female were separated, insuring that further male-female interactions such as re-mating attempts would not affect the estimate of fertility. Female body weight was measured prior to mating and male body weight both before and after mating to estimate the weight of the transferred ejaculate. After mating, while females were placed in 6cm petri dishes containing ad libitum (ca 40) black-eyed beans and left to lay eggs, males were scored for their locomotor activity.

Male locomotor activity was scored ca. 30 minutes following the single monogamous mating event. 20 males per replicate line per temperature were placed in groups of four in 6cm petri dishes on a heating plate that maintained the original assay temperature (29 or 36°C). After ten minutes of acclimation, each dish was observed every 30 seconds for ten minutes and considered active if one or more of the four individuals were moving (see: Berger et al 2016b).

Population fitness was measured at respective temperature as the average female offspring production in a group of 10 individuals with equal sex-ratio, placed together in a large petri-dish (6cm wide, 2cm deep) with ad libitum (ca. 200) black-eyed beans and left to interact for their lifetime.

### Statistics

#### Competitive LRS, fertility and population fitness

Our analyses used maximum likelihood estimation from general linear mixed effects models, assuming normally distributed data, implemented in the lme4 package (Bates *et al.* 2011) for R (R Core Team 2013). Evolution regime, assay environment, sex and their interactions were specified as fixed effects. Line identity (unique replicate line ID) crossed by environment and sex, as well as date of the assay, were specified as random effects. This thus ensured that the error variance between replicate lines within evolution regimes was used to evaluate statistical significance of all terms including evolution regime, and this general procedure was used in all analyses. For fertility, evolution regime, assay environment, as well as ejaculate weight were specified as fixed effects. Line identity crossed by temperature was specified as a random effect. Population fitness was analyzed in a model including the fixed effects of evolution regime, temperature and their interaction, and line identity crossed by temperature as a random effect. The normality of residuals was checked for all models.

#### Female and male weight, ejaculate weight and male activity

Body weight was analyzed using a liner mixed model assuming a Normal distribution. Evolution regime, temperature, sex and their interactions were specified as fixed effects and line identity and line crossed by temperature as random effects. Ejaculate weight was analyzed using the same model structure but for the main effect of sex. Finally, activity was analyzed using the same model structure as ejaculate weight but assuming a Binomial distribution for the response.

#### Cost of mating interactions

The change in per female offspring production (B) between monogamous fertility assays and population assays should capture the effect of mating interactions on female viability and reproduction. This change was calculated in relative terms as: 1 - B_population_ / B_fertility_.

To estimate B_population_ and B_fertility_, we ran a Bayesian mixed effects model implementing Markov Chain Monte Carlo simulations using the MCMCglmm package (Hadfield 2010) for R. Offspring count was the normally distributed response variable, with temperature, evolution regime, type of assay (fertility or population fitness) and their interactions as fixed effects. Line identity crossed by temperature and assay type were specified as random effects. Output of the model can be found in Supplementary table 8. Default weak priors were used (Variance initiated at 1 and belief set to 0.002 for all random effects) and the number of iterations was set to 1 100 000, of which the first 100 000 were used for burn-in and later discarded. We stored every 1000^th^ simulation, resulting in 1000 uncorrelated posterior estimates upon which we could calculated Bayesian 95% credible intervals for all parameter estimates and P-values for all comparisons.

We extracted posterior distributions for the mean offspring count (B) of each assay type (fertility or population fitness) by evolution regime and temperature combination. We then used these posterior distribution to estimate the cost of mating (1 - B_population_ / B_monogamy_) for each evolution regime and temperature combination. To assess if the cost of mating differed between two evolution regimes or temperatures (or combinations of those factors), we calculated the number of times the difference between posterior estimates of two groups being compared overlapped zero, with ≤ 2.5% cases implying statistical significance given a two-sided hypothesis. All the posterior distributions are reported in Table 10 with posterior mode and 95% credible intervals.

## Results

### Individual lifetime reproductive success of males and females

When measured in their ancestral environment (29°C on black eyed beans), evolution regimes differed in competitive lifetime reproductive success (LRS) (*χ*^2^_2_ =8.55, p=0.014, Fig.1). A post-hoc test indicated that the Polygamy regime was responsible for this effect (Tukey’s post-hoc, Male-limited – Monogamy: p=0.56, Polygamy – Monogamy: p=0.13, Polygamy – Male-limited: p=0.013). This suggests that sexual selection (present in both the Polygamy and Male-limited evolution regime), and male-female coevolution (present only in the Polygamy regime), are both important to maintain high individual LRS in a competitive context.

**Figure 1.**
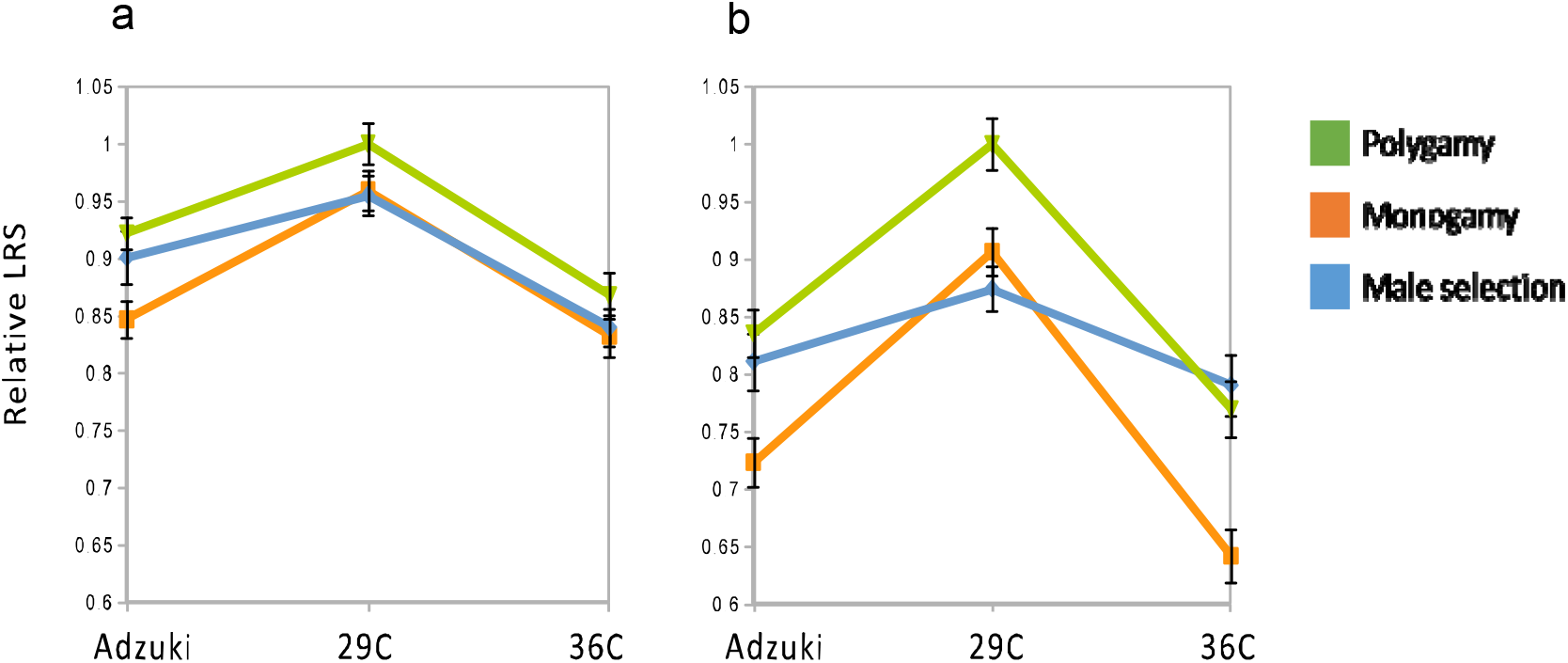
Sex-specific lifetime reproductive success (LRS) in each of the three assay environments in the three evolution regimes. Female (a) and male (b) LRS was standardized separately by mean LRS of the Polygamy regime at 29°C. Error bars represent one standard error.

The sexes responded differently to environmental stress (*χ*^2^_2_ =15.7, p<0.001, Table 1.a): while environmental stress (elevated temperature or new host plant) reduced LRS of both sexes, males were generally more affected than females (Fig. 1). A significant three-way interaction including sex, assay environment and evolution regime suggested that the regimes responded differently to environmental stress, and that this difference was sex-specific (*χ*^2^4=11.8 p=0.019, Table 1.a). To examine this further, we ran separate models for the sexes. This showed that the evolution regime by environment interaction was not found in females (*χ*^2^_4_=1.35, p=0.85, Table 1.b) but present in males (*χ*^2^_4_=22.7, p<0.001, Table 1.c). This is explained by the fact that males from the Male-limited evolution regime were overall relatively less affected by temperature or host plant than males from the other two regimes (Fig. 1b). This suggests that when sexual selection acted alone, alleles conferring environmental robustness were enriched relative to the other regimes that applied fecundity selection. Moreover, the increased robustness to elevated temperature in the male-limited regime was limited to males (Fig. 1). In females, there was no significant overall effect of evolution regime on competitive LRS (Table 1.b), but when comparing only the Monogamy and Polygamy evolution regimes a significant difference was found (Tukey’s post-hoc, p=0.033, visible on figure 1.a), suggesting that sexual selection and male female-coevolution increased female reproductive success as compared to fecundity selection.

**Table 1.** (a) Anova table for a general linear mixed-effect model of competitive lifetime reproductive success, showing the effect of sex, assay environment, evolution regime and their interactions. Tables (b) and (c) respectively show the analysis for females and males separately. P-values were calculated using type III sums of squares.

| Fixed Effect | $\chi^2$ | df | P-value |
| --- | --- | --- | --- |
| Evolution regime | 1.71 | 2 | 0.43 |
| Environment | 15.7 | 2 | <0.001 |
| Sex | 0.45 | 1 | 0.5 |
| Environment: Evolution regime | 1.85 | 4 | 0.76 |
| Sex : Evolution regime | 0.67 | 2 | 0.72 |
| Sex : Environment | 18 | 2 | <0.001 |
| Sex : Environment: Evolution regime | 11.8 | 4 | 0.02 |

b.
| Fixed Effect | $\chi^2$ | df | P-value |
| --- | --- | --- | --- |
| Evolution regime | 3.84 | 2 | 0.15 |
| Temperature | 14.6 | 2 | <0.001 |
| Temperature: Evolution regime | 1.35 | 4 | 0.85 |

| Fixed Effect | $\chi^2$ | df | P-value |
| --- | --- | --- | --- |
| Evolution regime | 5.2 | 2 | 0.07 |
| Temperature | 85.2 | 2 | <0.001 |
| Temperature: Evolution regime | 22.7 | 4 | <0.001 |

Next we explored the consequences of evolution under alternative mating regimes and the observed sex-specific temperature tolerance for fertility and population fitness at benign and elevated temperature.

### Fertility of male and female mating pairs

Fertility was measured as the lifetime offspring production of females following a single monogamous mating with a conspecific male. Evolution regimes differed in terms of fertility (*χ*^2^_2_=8.91, p=0.012, Supplementary table 1), with the Male-limited evolution regime having the highest fertility (Fig. 2). A post-hoc test indicated that the difference between Male-limited selection and Monogamy was driving this effect (Tukey’s post-hoc, Male-limited – Monogamy: p=0.008, Polygamy – Monogamy: p=0.32, Polygamy – Male-limited: p=0.27). This result thus provides evidence that Male-limited sexual selection did not result only in phenotypes that are successful in competitive settings, but also generated phenotypes with the highest fitness in monogamous settings. Being exposed to elevated temperature decreased fertility across all regimes but there was no significant interaction between regime and temperature (Supplementary table 1, Fig. 2).

**Figure 2.**
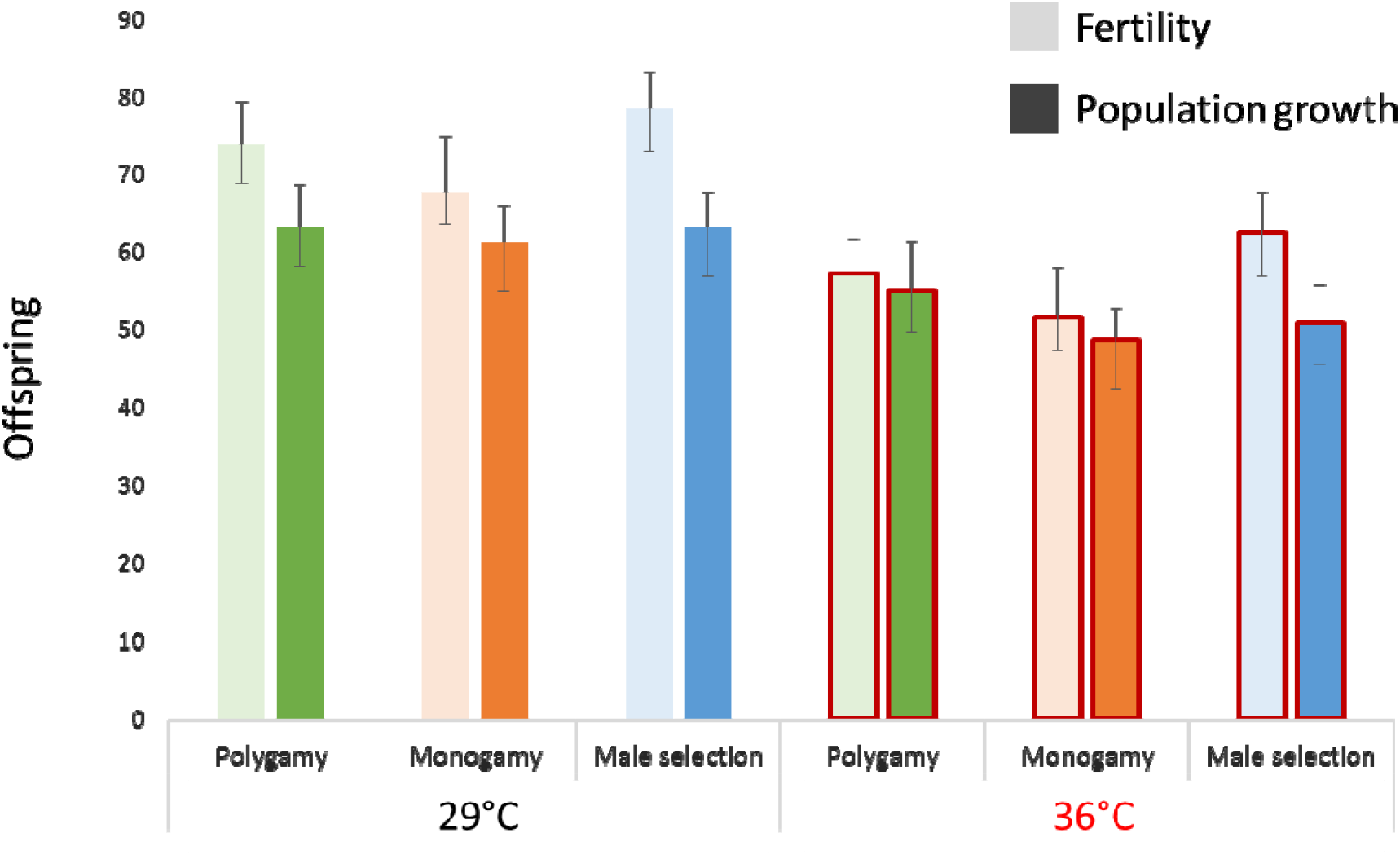
Fertility and population fitness at benign and elevated temperature in the three evolution regimes. Fertility (light bars) and population fitness (dark bars) was estimated by MCMC simulations and error bars represent 95% Bayesian credible intervals.

To provide insights into the differences in fertility across evolution regimes we performed an additional analysis on fertility where we added the three covariates to the model: female weight, male weight and male ejaculate weight (see: Supplementary figure 4). Females from the Polygamy regime are heavier than other females at both temperatures, which may explain the high fertility of Polygamy lines compared to Monogamy lines. However, females from the Male-limited evolution regime had similar weight as females from the Monogamy regime, yet the Male-limited regime exhibits the highest fertility at both temperatures (Fig. 2). This suggests that while the high fertility of the Polygamy regime relative to the monogamous regime may be explained by female size, this explanation does not account for the high fertility of the Male-limited regime. Ejaculate weight and male size showed no significant effects on fertility (Fig. S5).

### Population fitness

Population fitness was measured as the offspring production per female in a small population of ten individuals with equal sex ratio. Even though the Male-limited regime had highest fertility, population fitness tended to be highest in the Polygamy regime, and the significant difference between the Monogamy and Male-limited regime seen for the analysis of fertility was no longer apparent for population fitness (Fig. 2). Indeed, the only difference between evolution regimes close to being significant was that between Polygamy and Monogamy (Tukey’s post-hoc test, p=0.057), whereas Polygamy and Male-limited selection (p=0.40) and Monogamy and Male-limited selection (p=0.58) did not differ. The overall effect of evolution regime was marginally non-significant (*χ*^2^_2_= 5.24, p=0.073, Supplementary table 2). These results are consistent with population fitness being determined by a balance between “good genes” sexual selection and sexual conflict, both of which were presumably higher in the Male-limited regime than in the Monogamy regime, resulting in no obvious difference in population fitness between the two, despite clear differences in fertility (Fig 2). Moreover, the comparatively high population fitness of the Polygamy regime suggests that male-female coevolution had a positive effect on population fitness. Temperature stress significantly reduced population fitness in all evolution regimes (*χ*^2^_2_=115, <0.001 Supplementary table 2), with no statistically significant interaction between evolution regime and temperature (*χ*^2^_2_=3.41, p=0.18, Supplementary table 2, Fig. 2).

### Sexual conflict and the net cost of mating interactions

The cost of mating was estimated as one minus the ratio of offspring produced per female in the population setting relative to the fertility of monogamous pairs, thus giving the proportion of offspring lost per female due to socio-sexual interactions. This cost was greater than 0 in all 1 000 Bayesian simulations, indicating substantial costs of mating to population fitness (Fig. 3). There were, however, differences in the cost of mating across both temperatures and evolution regimes (Fig. 3). According to prediction, the Male-limited regime incurred a greater cost than the Polygamy regime averaged across temperatures (in 995 out of 1 000 simulations, two-sided Bayesian P = 0.01, Figure 3). This result thus directly demonstrates that experimental evolution of male adaptations under sexual selection without female coevolution confers strong costs at the population level. There was also a tendency for the cost to be greater in the Male-limited regime compared to the Monogamy regime, as expected if males evolving under sexual selection are more harmful to females than monogamous males, but the difference was marginally non-significant (959 out of 1 000 simulations, two-sided Bayesian P = 0.08).

**Figure 3.**
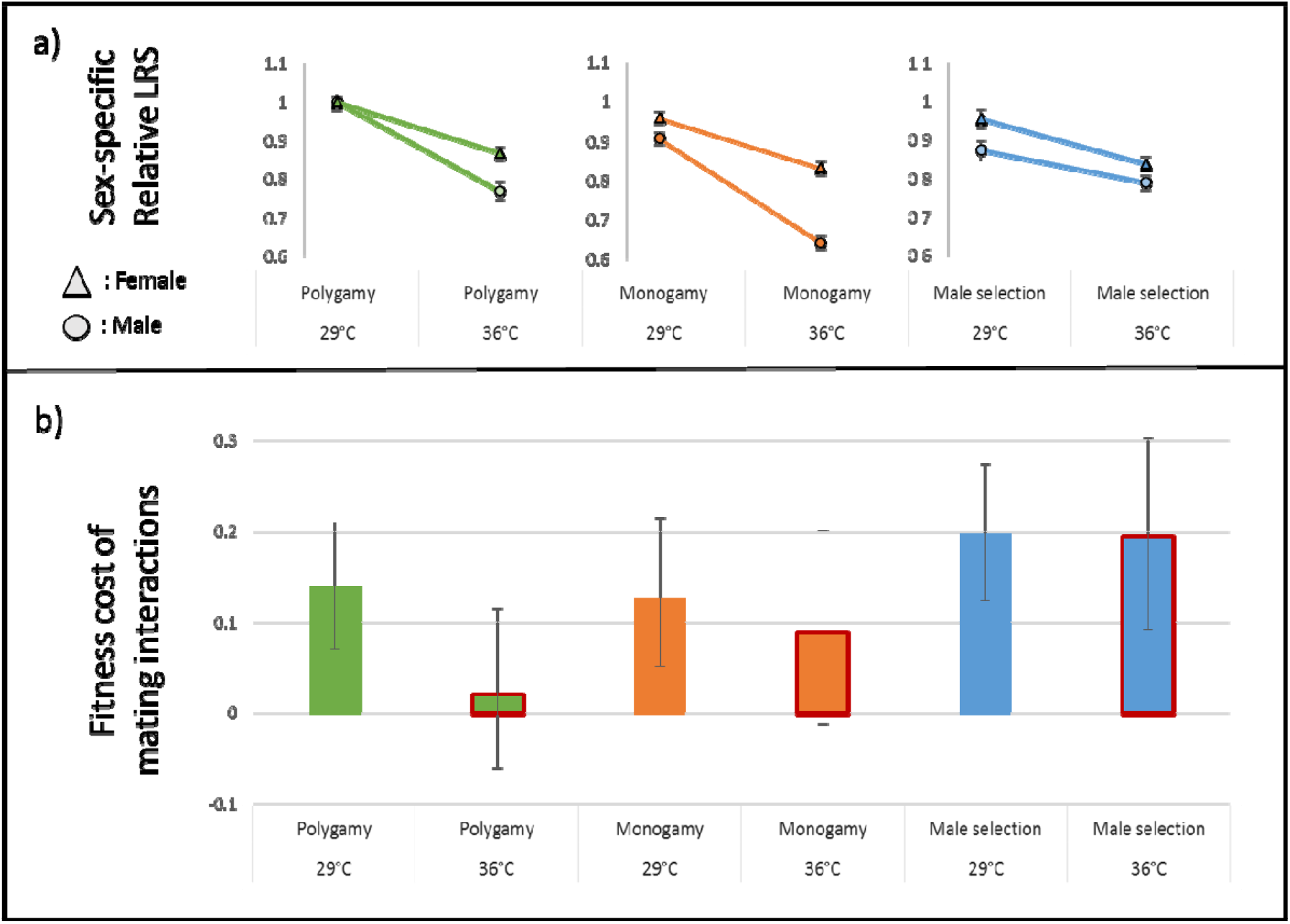
Sex-specific sensitivity to elevated temperature and associated cost of mating interactions for the three evolution regimes. a) Sex-specific relative LRS is presented separately for each evolution regime to emphasize sex-specific temperature-sensitivity in each regime. Error bars represent standard errors. b) The cost of mating interactions was calculated as *1-(population fitness/fertility)* and represents the relative drop in per capita offspring production between a single monogamous mating and a polygamous group setting. Error bars represent 95% Bayesian credible intervals.

There was an overall tendency for elevated temperature to reduce the cost of mating (942 out of 1 000 simulations, Fig. 3). However, the effect of temperature was not consistent across regimes. In the Polygamy and Monogamy regimes, the cost of mating appeared to be reduced at elevated temperature, although only markedly so for Polygamy (966 out of 1 000 simulations, two-sided Bayesian P-value = 0.07). In contrast, the cost of mating interactions in the Male-limited regime remained high and constant. The elevated temperature magnified the difference in the cost of mating between Polygamous and Male-limited evolution regime: 15% cost vs 19% cost at 29°C, p=0.13; 0.5% cost vs 19% cost at 36°C, p=0.02. In Figure 3, we present this result in parallel with the sex-specific temperature-sensitivity of LRS in order to explore the link between sex-specific LRS and population dynamics and the impact of temperature stress on that relationship (see also Fig. 1).

In an attempt to unveil the mechanisms and phenotypes mediating the costs of mating, we revisited our data on individual male and female traits measured in the fertility assays. While male and female body weight both increased at elevated temperature (*χ*^2^_2_=35.2, p<0.001, Supplementary table 3.a, Supplementary figure 5), we could not reveal any interactions between sex, temperature and evolution regime, suggesting that sex-specific responses in body weight do not explain putative variation in the cost of mating. Similarly, ejaculate weight was not affected by either temperature or evolution regime (Supplementary table 3.b), which gives no support for the idea that the amount of seminal fluid proteins, nutrition, or toxic compounds in the ejaculate were central to generating the observed variation. We note, however, that the relative composition of these components may still have influenced mating costs. Finally, male locomotor activity, which gives an indication of male harassment in seed beetles (e.g. Berger *et al.* 2016a), decreased at elevated temperature (*χ*^2^_2_=5.80, p=0.016, Supplementary table 3.c, Supplementary figure 6), but this decrease was similar in all evolution regimes. Hence, while possibly explaining parts of the reduced cost of mating interactions in the monogamous and polygamous mating regime, something additional and unidentified must in that case explain the maintained mating cost in the Male-limited regime.

## Discussion

The impact of sexual selection on adaptation results from a balance between the benefits of good genes effects and costs of sexual conflict. If these processes are affected by environmental change, this balance could be offset in maladapted populations. However, exactly how these effects will be manifested in novel environments remains unknown because of the lack of empirical data, and here we therefore tried to explore these dynamics. We used sex-limited experimental evolution to disentangle the respective contributions of sexual selection, fecundity selection and male-female coevolution, to individual-level and population-level fitness. We then contrasted these effects in well-adapted populations raised at ancestral conditions, and maladapted populations raised at elevated temperature. Our study demonstrates how sex-specific selection and maladaptation can affect the link between individual-level adaptation and population viability in polygamous species.

### Good genes, environmental robustness, and genotype-by-environment interactions

Whether sexual selection generally results in good genes effects (Tomkins et al. 2004; Bonduriansky and Chenoweth 2009), and whether such effects persist in novel environments (Bussiere et al. 2008; Radwan 2008; Kolluru et al. 2014), remains a matter of considerable debate. In our experiment, the Male-limited evolution regime showed the highest fertility of all three evolution regimes, and a female reproductive success similar to the Monogamy regime (Fig. 2), suggesting that sexual selection on males can indeed increase female fitness components in *C. maculatus*. In this species, energetically costly interference and scramble competition is intense (Savalli and Fox 1999; Maklakov and Arnqvist 2009), making it likely that males of high quality that carry “good genes” are favored by sexual selection (Whitlock and Agrawal 2009). In addition to good genes effects, there are other, non-mutually exclusive, mechanisms that may have contributed to the generally high female reproductive output in both Polygamy and Male-limited lines. For example, manipulation of female physiology by males during mating (for example through the medium of seminal fluid proteins), could have evolved or been enhanced by male competition under sexual selection. Indeed, such effects are known from other model systems such as fruit flies and nematodes (e.g: Chapman et al. 1995; Gems and Riddle 1996). We also cannot rule out that sexual selection acted on traits that are not condition dependent per se but on indicators of alternative genotypes for which mate preferences are dictated by the complementarity of the mating partners, such as has been reported for MHC-linked traits (Wedekind 1994; Ruff et al. 2012).

Males from the male-limited evolution regime showed high resilience to both host plant and temperature stress, suggesting that sexual selection on males lead to environmentally robust phenotypes that perform well across environments rather than locally adapted specialists (see also: Parrett and Knell 2018). Interestingly, males from the polygamous regime (that also applied sexual selection) did not show the same environmental robustness. This suggests that a balance between fecundity selection and sexual selection is important in shaping sex-specificity in environmental robustness, and may be central in maintaining alternative alleles encoding this trait in *C. maculatus* populations. Such alleles are likely to contribute to adaptation in new environments, supporting the idea that opposing forces of natural and sexual selection can maintain genetic variation that may fuel adaptive responses to environmental change (Radwan et al. 2016). We cannot rule out sexual selection also acting on traits that are not condition dependent but rather indicators of variability or complementarity of the reproduction partner such as MHC-linked traits (Ruff et al. 2012; Wedekind 1994).

### Sexual conflict and population demography upon environmental change

Despite leading to genetic increases in female fitness components, we also saw that sexual selection can favor individual male strategies that bear costs at the population level. We observed a cost of mating in all evolution regimes at the ancestral temperature (Fig.3). This is a clear sign of IeSC, which is not surprising in this species, well known for costs of multiple mating in females and sexually antagonistic coevolution involving male and female genitalia (Crudgington and Siva-Jothy 2000; Edvardsson and Tregenza 2005; Rönn *et al.* 2007; Gay *et al.* 2011; Dougherty et al. 2017). Our study also demonstrates that IeSC evolves to become magnified under sexual selection, especially when female counter-adaptations are constrained from mitigating the harm incurred by male mating strategies (Rice 1996), signified by the higher cost of mating in Male-limited relative to both Monogamous and Polygamous lines. This result is consistent with theoretical models suggesting that sexual selection can lead to the evolution of male traits that are harmful to females and thereby can contribute to population extinction (Kokko and Brooks 2003; Rankin *et al.* 2011). This view is also in agreement with recent evidence from the fossil record showing that lineages of ostracods with higher sexual dimorphism, as a correlate of the strength of sexual selection and conflict, have higher rates of extinction (Martins *et al.* 2018).

The potential impact of IeSC on population viability has been discussed (Parker 1979; Thornhill and Alcock 1983; Clutton-Brock and Parker 1995; Arnqvist and Rowe 2005; Parker 2006), and recent empirical studies have explored its effects across variable ecological conditions (e.g. Rowe and Arnqvist 2002; Sakurai and Kasuya 2008; den Hollander and Gwynne 2009; Gay et al. 2009; Berger et al. 2012; Arbuthnott et al. 2014; Takahashi et al. 2014; Gomez-Llano 2018; Skwierzynska et al. 2018). However, making predictions about the extent and change in IeSC and its consequential impact on population demography upon environmental change is complicated by the inherent unpredictability of environmental change itself. For example, it has been argued that IeSC may be reduced under low population density (Arnqvist 1992; Gerber and Kokko 2016), as can be expected in declining populations suffering from maladaptation to local environmental conditions. In this case, IeSC would be relaxed and the population relieved of the sexual conflict load. However, depending on the context driving population decline, it remains uncertain whether general declines in numbers of breeding pairs will result in lower densities at mating sites, especially if the drivers of population decline are factors like degradation and fragmentation of suitable habitat for breeding, which may instead result in higher densities of reproducing adults. Here, therefore, we raised one possible heuristic scenario that could generate predictable changes in IeSC and its demographic cost upon environmental change. If environmental stress affects one sex more than the other (for example because of different resource use: Maklakov et al. 2008; Zajitschek and Connallon 2017), IeSC could either be intensified or reduced depending on which sex is the most sensitive to the change in ecological conditions (Clutton-Brock and Parker 1995; Rankin et al. 2011; Iglesias-Carrasco *et al.* 2018). Interestingly, the greater female-bias in environmental robustness found in the Polygamy and Monogamy regime, relative to the Male-limited regime where fecundity selection was removed, suggests that the balance between natural and sexual selection may shape sex-specific environmental robustness. As follows from our hypothesis, these sex-differences in environmental robustness were accompanied by parallel changes in the cost of mating at elevated temperature, with lowered costs observed in the Monogamy and Polygamy regime, (although this difference was significant only in the Polygamy regime) but maintained costs in the Male-limited regime (Fig. 3). This thus raises the possibility that sex-specific selection can lead to sex-specificity in environmental robustness which in turn can modulate the cost of sexual conflict. While this principle may be general, it remains to be explored how great its effect is in natural populations facing changing ecological settings, and how predictable sex-differences in environmental robustness are in naturally variable environments. Both these aspects, along with the fact that multiple mating also may provide females with benefits that are likely affected by the condition of the male (Arnqvist and Nilsson 2000), will need to be understood in order to form a better understanding of the demographic impact of the explored dynamics.

### Traits underlying male-induced harm

One way to better understand how the relationship between sex-specific environmental robustness and IeSC can be generalized to other animals may be to investigate the traits that mediate sexual conflict. Body weight is a good candidate trait, as it often reflects condition and can be a mediator of sexual conflict (Clutton-Brock and Parker 1995). Indeed, sexual size dimorphism is likely to be related to the ability of males inflicting harm, or females’ ability to cope with harmful male behaviors, during mating (Arnqvist and Rowe 2005). In *D. melanogaster* for example, the relative body weight of the sexes determines the impact of sexual conflict on female fecundity (Pitnick and García-González 2002; Friberg and Arnqvist 2003). Hence, because body size is condition dependent in many species, sex-specific sensitivity to stress may alter the degree of sexual size dimorphism (Stillwell et al. 2010), and thereby the intensity of sexual conflict, in predictable ways. In the present study however, body weight did not explain the cost of mating interactions. The case of body weight could be extended to other condition dependent traits that are involved in sexual conflict. Male locomotor activity, which is a likely cause for reduced female offspring production in many species (Arnqvist and Rowe 2005), and in *C. maculatus* (Gay *et al.* 2009; Berger *et al.* 2016a), was lower at elevated temperature. However, male activity did not differ significantly between evolution regimes, suggesting that it was not the trait responsible for the maintained cost of mating at elevated temperature in the Male-limited regime. Finally, contrary to what has been observed in insect taxa like *Drosophila melanogaster* (Lung *et al.* 2002; Wigby and Chapman 2005; Mueller *et al.* 2007), we found no support for ejaculate toxicity mediating IeSC to the extent that ejaculate weight did not vary across evolution regimes. We do note that this does not rule out the possibility that the composition of the ejaculate (e.g. Goenaga *et al.* 2015) may have played a role in generating variation in both fertility and IeSC across temperatures and evolution regimes.

In summary our study points to multiple facets by which sexual selection can either contribute to evolutionary rescue or extinction of maladapted populations. Our results highlight that these effects can be interdependent because sexual selection on males can (i) elevate fertility of females via good genes effects, but also (ii) intensify sexual conflict, and (iii) involve loci with environment-specific effects, affecting direct as well as genetic responses in novel environments, and finally (iv) affect sex-specific sensitivity to environmental stress, which in turn may modulate the intensity of sexual conflict in maladapted populations facing novel ecological challenges.

## Acknowledgements

This work was funded by a grant from the Swedish Research council VR to DB (grant no. 2015-05233) and GA (621-2014-4523), and a grant from the European Research Council (ERC) to GA (GENCON AdG-294333). We are thankful to J. Liljestrand-Rönn for invaluable help during experiments, and to B. Stenerlöw for providing access to the radiation source.

## Data Archiving

Data for this study are available at: to be completed after manuscript is accepted for publication.

---

**Supplementary table 1.** Anova table for a general linear mixed-effect model of fertility, showing the effect of assay temperature, evolution regime and ejaculate weight. P-values were calculated using type II sums of squares.

| Fixed Effect | $\chi^2$ | df | P-value |
| --- | --- | --- | --- |
| Evolution regime | 8.91 | 2 | 0.01 |
| Temperature | 39.5 | 1 | <0.001 |
| Ejaculate weight | 0.07 | 1 | 0.79 |

**Supplementary table 2.** Anova table for a general linear mixed-effect model on population fitness, showing the effect of assay temperature, evolution regime and their interactions.

| Fixed Effect | $\chi^2$ | df | P-value |
| --- | --- | --- | --- |
| Evolution regime | 5.24 | 2 | 0.073 |
| Temperature | 115 | 1 | <0.001 |
| Temperature: Evolution regime | 3.41 | 2 | 0.18 |

**Supplementary table 3.a.** Anova table for a general linear mixed-effect model on body weight showing the effect of assay temperature, evolution regime, sex and their interactions.

| Fixed Effect | $\chi^2$ | df | P-value |
| --- | --- | --- | --- |
| Evolution regime | 4.50 | 2 | 0.11 |
| Temperature | 35.2 | 1 | <0.001 |
| Sex | 1117 | 1 | <0.001 |
| Temperature: Evolution regime | 0.784 | 2 | 0.68 |
| Sex : Evolution regime | 4.26 | 2 | 0.12 |
| Sex : Temperature | 2.15 | 1 | 0.14 |
| Sex : Temperature: Evolution regime | 0.0073 | 2 | 0.99 |

**Supplementary table 3.b.** Anova table for a general linear mixed-effect model on male ejaculate weight, showing the effect of assay temperature, evolution regime, and their interactions.

| **Fixed Effect** | **$\chi^2$** | **df** | **P-value** |
| --- | --- | --- | --- |
| Evolution regime | 1.13 | 2 | 0.57 |
| Temperature | 0.083 | 1 | 0.77 |
| Temperature: Evolution regime | 2.72 | 2 | 0.26 |

**Supplementary table 3.c.** Anova table for a generalized linear mixed-effect model on male activity, assuming Binomial errors, showing the effect of assay temperature, evolution regime and their interactions.

| Fixed Effect | $\chi^2$ | df | P-value |
| --- | --- | --- | --- |
| Evolution regime | 2.01 | 2 | 0.37 |
| Temperature | 5.80 | 1 | 0.016 |
| Temperature: Evolution regime | 0.151 | 2 | 0.93 |

**Supplementary figure 4.**
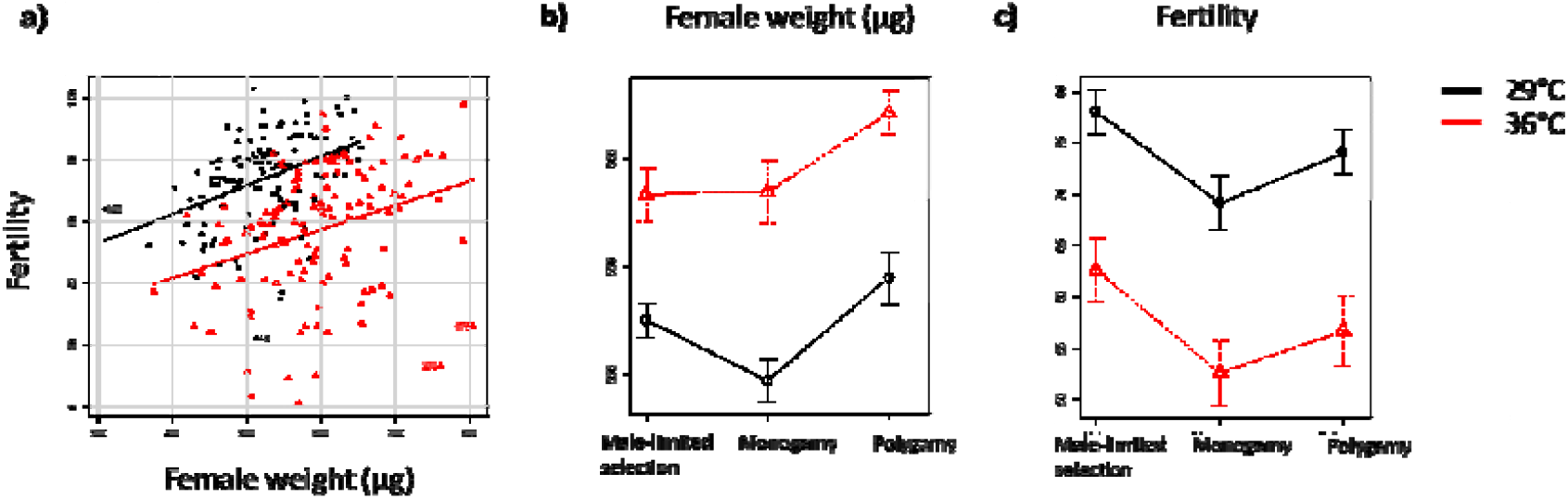
The relationship between female weight and fertility across temperatures and experimental evolution treatments.

**Supplementary figure 5.**
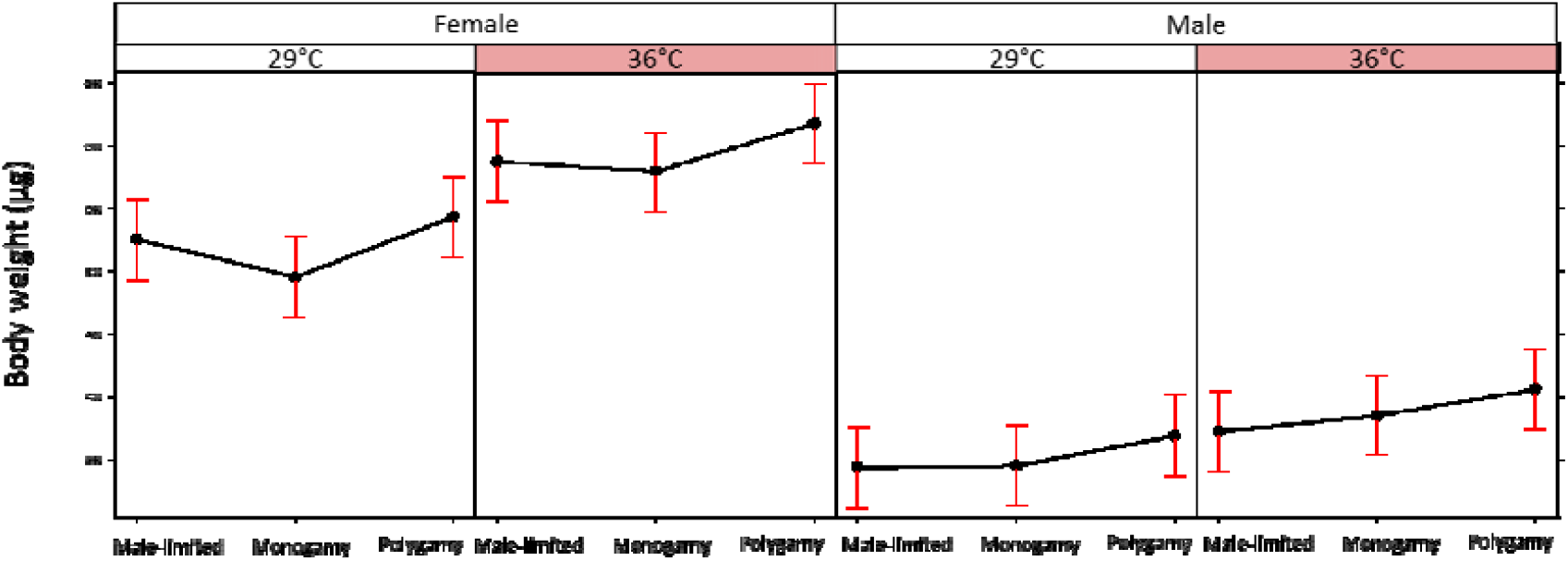
The effect of temperature and evolution regime on female and male weight.

**Supplementary figure 6.**
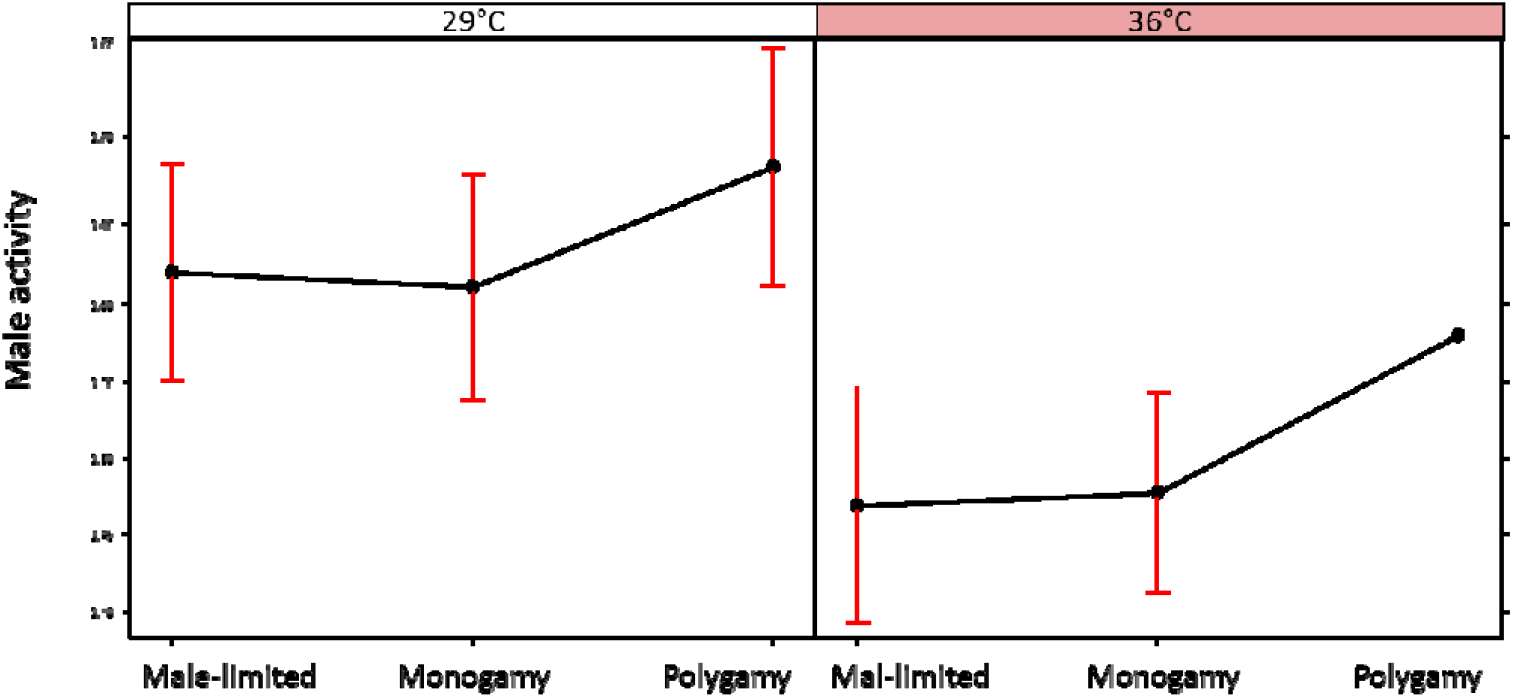
The effect of temperature and evolution regime on male activity.

**Supplementary figure 7.**
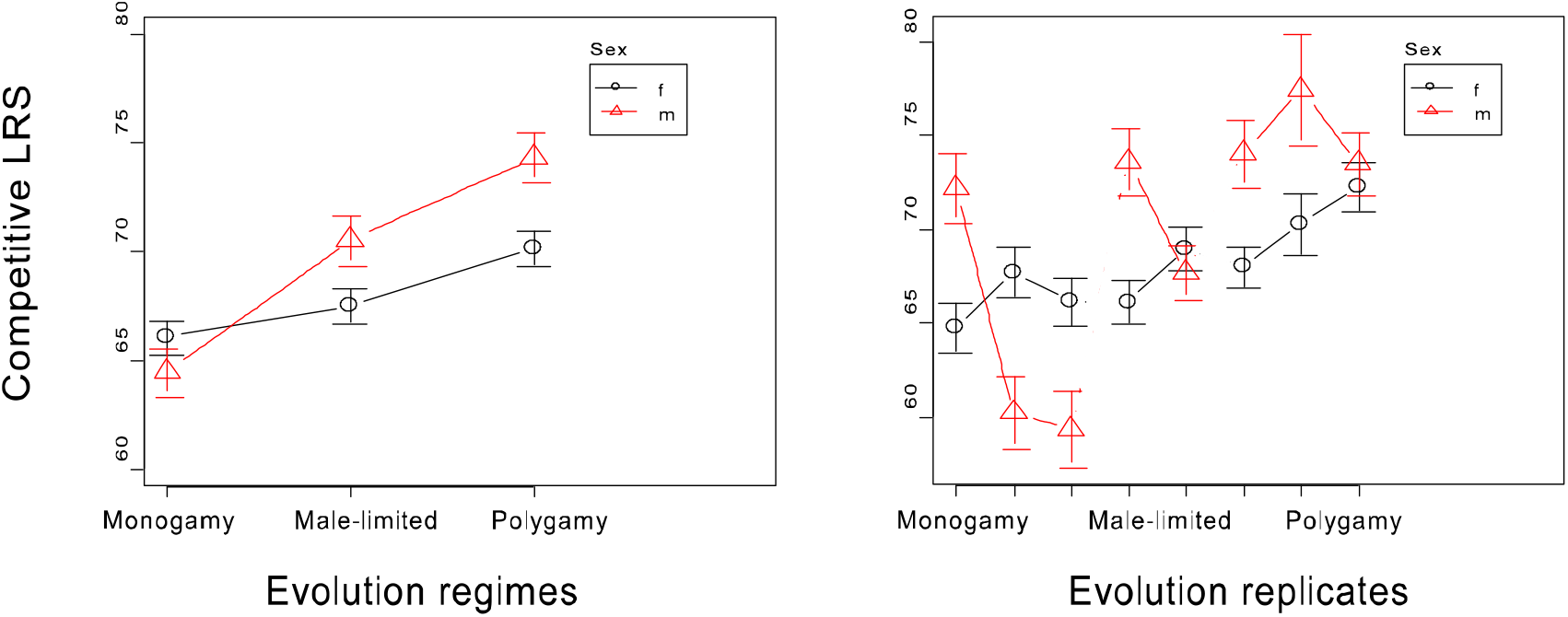
Mean absolute LRS for each evolution regime (left pannel) and each replicate line of the evolution regimes (right pannel), for females (circles) and males (triangles).

**Supplementary table 8.** Output of the MCMCglmm model used to estimate the cost of mating interactions. The response variable was offspring count; evolution regime, temperature and type of assay (population fitness or fertility assay) as well as their interactions were specified as fixed effects. Evolution replicate, temperature by replicate and assay type by replicate were specified as random effects.

| **Fixed Effect** | **Posterior mean** | **95% CI** | **P<sub>MCMC</sub>** |
| --- | --- | --- | --- |
| Intercept | 62.45 | 56.03/68.27 | <0.001 |
| Assay type: Fertility | 15.73 | 9.38/22.02 | <0.001 |
| Selection: Monogamy | -1.89 | -12.17/7.06 | 0.67 |
| Selection: Polygamy | 0.98 | -7.52/10.97 | 0.81 |
| Temperature: 36C | -11.7 | -19.12/-3.99 | 0.01 |
| Fertility by Monogamy | -7.19 | -16.64/1.87 | 0.13 |
| Fertility by Polygamy | -5.15 | -15.31/3.31 | 0.25 |
| Fertility by 36C | -3.67 | -11.85/4.76 | 0.38 |
| Monogamy by 36C | -0.59 | -12.13/9.98 | 0.92 |
| Polygamy by 36C | 3.7 | -7.10/15.51 | 0.46 |
| Fertility by Monogamy by 36C | -0.71 | -11.73/12.52 | 0.9 |
| Fertility by polygamy by 36C | -5.65 | -16.59/6.58 | 0.34 |
| **Random effects** |  |  |  |
| Evolution Replicate | 5.92 | 0.0031/21.0 |  |
| Temperature by Replicate | 7.76 | 0.0034/29.1 |  |
| Assay type by Replicate | 1.75 | 0.0035/8.0 |  |
| Residuals | 182.2 | 160.5/207.5 |  |

**Supplementary table 9.**
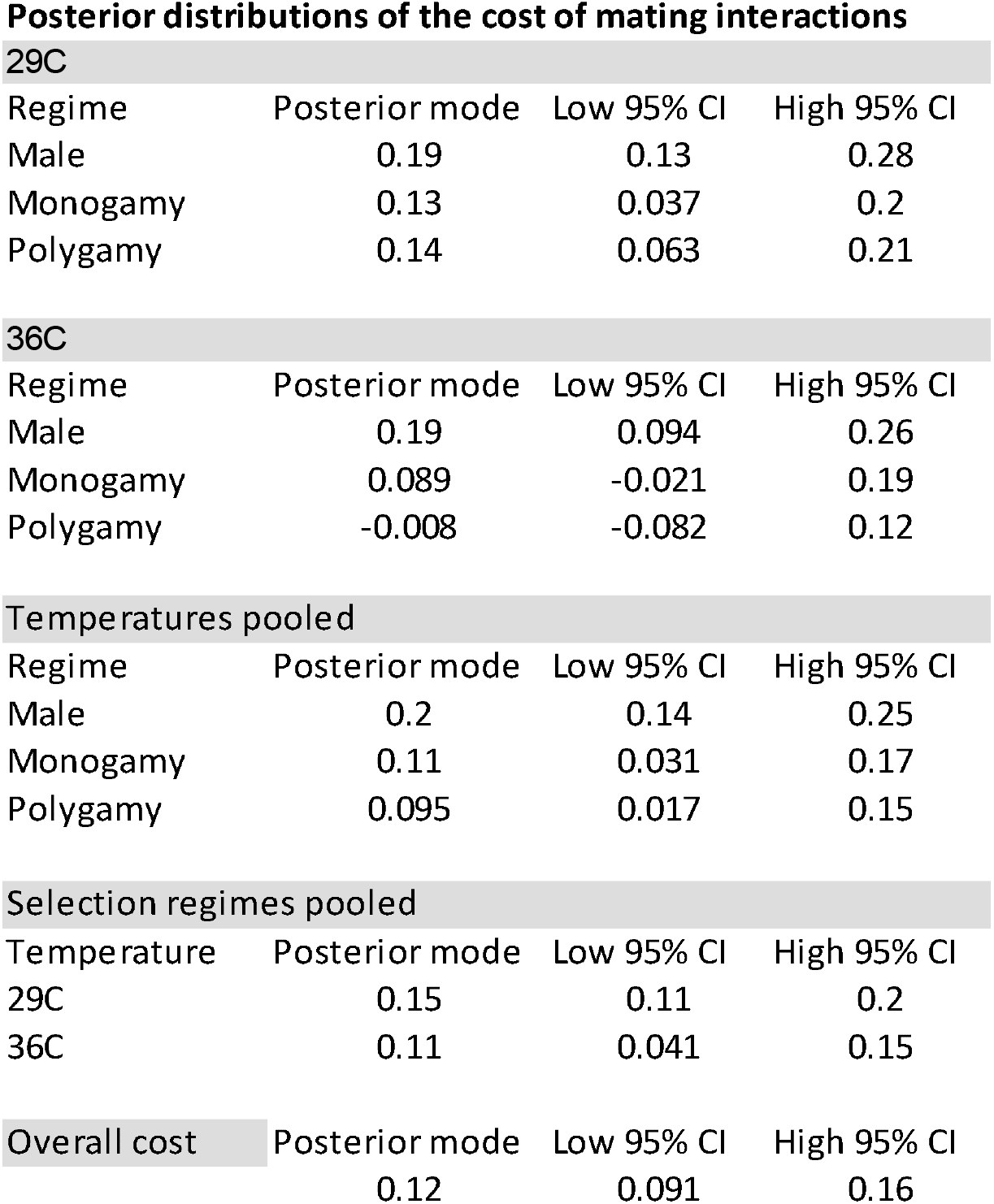
Posterior distributions extracted from an MCMCglmm model for the cost of mating calculated as 1 - B_population_ / B_monogamy_ were B represents mean offsprring production for respectively population assays and fertility assayed in a monogamous setting. Costs of mating are given for each evolution regime and temperature combination, as well as cost per evolution regime with temperatures pooled, per temperature with regimes pooled and finally overall cost.

| Regime | Posterior mode | Low 95% CI | High 95% CI |
| --- | --- | --- | --- |
| Male | 0.19 | 0.13 | 0.28 |
| Monogamy | 0.13 | 0.037 | 0.2 |
| Polygamy | 0.14 | 0.063 | 0.21 |

36C
| Regime | Posterior mode | Low 95% CI | High 95% CI |
| --- | --- | --- | --- |
| Male | 0.19 | 0.094 | 0.26 |
| Monogamy | 0.089 | -0.021 | 0.19 |
| Polygamy | -0.008 | -0.082 | 0.12 |

Temperatures pooled
| Regime | Posterior mode | Low 95% CI | High 95% CI |
| --- | --- | --- | --- |
| Male | 0.2 | 0.14 | 0.25 |
| Monogamy | 0.11 | 0.031 | 0.17 |
| Polygamy | 0.095 | 0.017 | 0.15 |

Selection regimes pooled
| Temperature | Posterior mode | Low 95% CI | High 95% CI |
| --- | --- | --- | --- |
| 29C | 0.15 | 0.11 | 0.2 |
| 36C | 0.11 | 0.041 | 0.15 |

| Posterior mode | Low 95% CI | High 95% CI |
| --- | --- | --- |
| 0.12 | 0.091 | 0.16 |

